# A statistical approach for tracking clonal dynamics in cancer using longitudinal next-generation sequencing data

**DOI:** 10.1101/2020.01.20.913236

**Authors:** Dimitrios V. Vavoulis, Anthony Cutts, Jenny C. Taylor, Anna Schuh

## Abstract

Tumours are composed of genotypically and phenotypically distinct cancer cell populations (*clones*), which are subject to a process of Darwinian evolution in response to changes in their local micro-environment, such as drug treatment. In a cancer patient, this process of continuous adaptation can be studied through next-generation sequencing of multiple tumour samples combined with appropriate bioinformatics and statistical methodologies. One family of statistical methods for clonal deconvolution seeks to identify groups of mutations and estimate the prevalence of each group in the tumour, while taking into account its purity and copy number profile. These methods have been used in the analysis of cross-sectional data, as well as for longitudinal data by discarding information on the timing of sample collection. Two key questions are how (in the case of longitudinal data) can we incorporate such information in our analyses and if there is any benefit in doing so. Regarding the first question, we incorporated information on the temporal spacing of longitudinally collected samples into standard non-parametric approaches for clonal deconvolution by modelling the time dependence of the prevalence of each clone as a *Gaussian process*. This permitted reconstruction of the temporal profile of the abundance of each clone continuously from several sparsely collected samples and without any strong prior assumptions on the functional form of this profile. Regarding the second question, we tested various model configurations on a range of whole genome, whole exome and targeted sequencing data from patients with chronic lymphocytic leukaemia, on liquid biopsy data from a patient with melanoma and on synthetic data. We demonstrate that incorporating temporal information in our analysis improves model performance, as long as data of sufficient volume and complexity are available for estimating free model parameters. We expect that our approach will be useful in cases where collecting a relatively long sequence of tumour samples is feasible, as in the case of liquid cancers (e.g. leukaemia) and liquid biopsies. The statistical methodology presented in this paper is freely available at github.com/dvav/clonosGP.

## INTRODUCTION

It is well known that cancer cells undergo a process of Darwinian evolution in response to selective pressures in their local micro-environment, for example as a result of therapeutic intervention[1, 2]. This induces cell propagation and diversification during tumour growth, which result in a heterogeneous population of phylogenetically related, but genotypically and phenotypically distinct cancer cell populations, known as *clones*. Tumour heterogeneity is clinically important because it complicates the molecular profiling of tumours and enables the fittest cancer cells to escape treatment leading to relapse. Monitoring this process of continuous adaptation requires a detailed characterisation (through the use of next-generation sequencing, bioinformatics and statistical analysis) of the somatic aberrations harboured by the tumour at various time points over the course of the disease.

A major challenge in solving the problem of *clonal deconvolution* using bulk sequencing data is the fact that tumour heterogeneity is not directly observed, but rather inferred through the analysis of samples, each of which is a mixture of normal and cancer cells from various clones. Despite (or because of) this, clonal deconvolution has been the subject of much statistical innovation (see [3–6] for a review). Current statistical methodologies seek to identify the number of clones in a tumour, their somatic mutation content, prevalence and phylogenetic relations and they can be used for the analysis of cross-sectional data (obtained, for example, through multiple biopsies from the same patient) or longitudinal data after discarding any information on the timing of tissue sample collection[7–22].

In this paper, we pose the following two questions: a) how can we incorporate temporal spacing information in the analysis of sequentially collected samples (typically over several months or years) and b) is there any benefit in doing so? We begin with a standard Bayesian non-parametric model for clustering somatic mutations with similar observed frequencies, while simultaneously correcting for sample purity and local copy number variation. We extend this model by treating the cluster prevalences as functions of time, which follow a Gaussian process prior. The advantage of this approach is that we do not need to impose a particular functional form on the time dependence of cluster abundances, but only some general properties (e.g. smoothness, amplitude and time scale), which are estimated from the data. In return, we obtain a continuous reconstruction of the time course of each cluster during the course of the disease from a small number of sequentially collected samples. We test various model configurations on whole genome (WGS), whole exome (WES) and targeted sequencing (TGS) data from patients with chronic lymphocytic leukaemia (CLL; [23, 24]), on data from the liquid biopsy of a patient with melanoma[25] and on synthetic data, and we demonstrate that incorporating temporal information in our analysis can boost the performance of clonal deconvolution.

## METHODS

We present a series of models of increasing complexity starting with the statistical model for a single tumour sample.

### Model for a single tumour sample

We assume that a tumour has been sequenced at *N* bi-allelic genomic loci harbouring somatic mutations. For each locus *i*, we can calculate the observed *variant allele fraction* (VAF) as the ratio 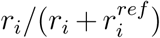, where *r*_*i*_ and 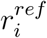 are the number of reads harbouring the alternative and reference alleles, respectively. The expected value *θ*_*i*_ of the VAF for mutation *i* is a function *f* of the *cancer cell fraction* (CCF), i.e. the fraction 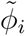 of cancer cells that harbour the mutation, 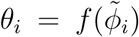. The population of cancer cells is partitioned in a finite, but unknown, number of clones, each harbouring a unique set of mutations. This implies that different mutations share the same CCF value, i.e. the mutation-specific fractions 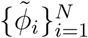 are not all distinct. We model this structure with a *Dirichlet Process* prior on 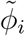 with concentration parameter *α* and a uniform base distribution 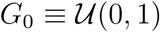[26, 27]. Using the stick-breaking representation of the Dirichlet Process, 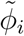 is modelled as an infinite mixture, as shown below:

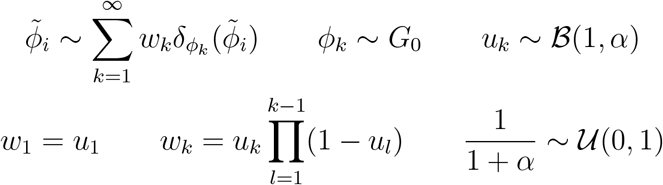

where 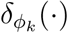 is the Dirac delta function centred at *ϕ*_*k*_ and 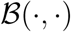 indicates a beta distribution. The uniform prior on the mean of the beta function (1 + *a*)^*−*1^ implies that the prior on the concentration parameter is *α* ∼ (1 + *α*)^*−*2^, which is similar to the standard exponential distribution, but with thicker tail. In practice, we truncate the above infinite sum at a value *K* larger than the maximum possible number of clones expected in the data (here taken equal to 20).

### Joint model for clonally-related tumor samples

The above model can be extended to multiple clonally-related samples by allowing the CCF variables to vary between samples[9, 28]. For *M* samples (and truncation *K*), we have:

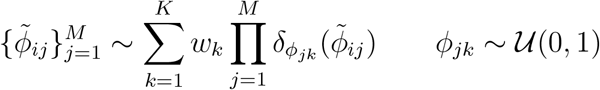

where the rest of the model remains the same as for the one-sample case. Effectively, we incorporate multiple samples in the model by allowing the cluster centres *ϕ*_*jk*_ to vary across samples. As a prelude to the next section, we note that the transformed variable *ψ*_*jk*_ = log *ϕ*_*jk*_ − log(1 *− ϕ*_*jk*_) follows a standard logistic distribution, *ψ*_*jk*_ ∼ Logistic(0, 1). Below, instead of the logistic distribution, we use a parametrised multivariate normal distribution, as explained in more detail in the next section.

### Single-output Gaussian Process model for longitudinal tumour samples

The above model does not take into account the temporal spacing of the *M* samples, in case these have been collected longitudinally. If such information is indeed available, it can be included in the model by treating the transformed CCF variables as functions of time, *ψ*_*k*_(*t*). On these functions, we impose a *Gaussian Process* prior[26, 29, 30]:

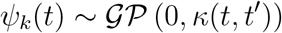

where the kernel function *κ*(*t, t*′) encodes the covariance of *ψ*_*k*_(*t*) at times *t* and *t*′. This non-parametric approach permits modelling the time-dependency of the transformed CCF variables without any strong prior assumptions on the functional form of this dependency. The above implies that if *M* samples have been collected at times *t*_1_ = 0, …, *t*_*j*_, …, *t*_*M*_ = 1, then the variables *ψ*_*jk*_ = *ψ*_*k*_(*t*_*j*_) follow a multivariate Normal distribution:

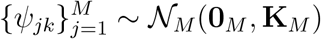

where **0**_*M*_ is the *M* -dimensional zero vector. The elements of the covariance matrix **K**_*M*_ = *{κ*(*t*_*j*_, *t*_*j′*_)}_*j,j′*_ encode the covariance between the values of *ψ*_*k*_(*t*) at all possible pairs of sampling times *t*_*j*_ and *t*_*j′*_.

We consider kernels of the form *κ*(*t, t*′) = *h*^2^*g*_*τ*_ (*t, t*′), where *h* is an amplitude parameter, while the function *g*_*τ*_ (*t, t*′), which is parametrised by an inverse squared time scale parameter *τ*, takes any of the following forms: a) exponential: 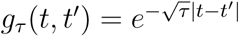, b) 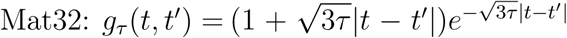, c) 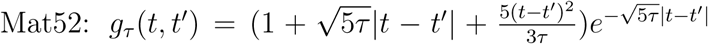 and d) exponentiated quadratic: 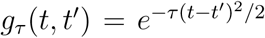. These four kernels are members of the Matern family of covariance functions ordered in terms of increasing smoothness[29]. Finally, we impose gamma priors on the amplitude and time scale parameters, 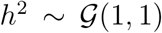 and 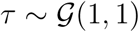.

### Multi-output Gaussian Process model for longitudinal tumour samples

In the above model, the cluster-specific scalar-valued functions *ψ*_*k*_(*t*) share the same Gaussian Process prior, but they are otherwise independent. We can directly model possible correlations between different clusters (i.e. different values of *k*) by assuming that the vector-valued function of time, 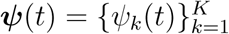, follows a Gaussian Process prior:

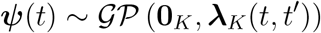

where ***λ***_*K*_(*t, t*′) is a matrix-valued kernel encoding the *K × K* covariance matrix between vectors ***ψ***(*t*) and ***ψ***(*t*′). Given *M* longitudinally observed samples, the above implies that the matrix of CCF values **Ψ**_*M×K*_ = *{ψ*_*jk*_}_*j,k*_ follows a multivariate Normal distribution of dimensionality *MK*:

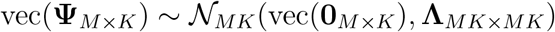

where the operator vec(*⋅*) vectorises its matrix argument by stacking its columns on top of each other, **0**_*M×K*_ is a matrix of zeros and **Λ**_*MK×MK*_ is a positive semi-definite block matrix encoding the covariance between *ψ*_*jk*_ and *ψ*_*j′k*′_.

Assuming that the above kernel is *separable*[31], we can write the factorisation ***λ***_*K*_(*t, t*′) = *g*_*τ*_ (*t, t*′)**Σ**_*K*_, where *g*_*τ*_ (*t, t*′) is the same as in the previous section. **Σ**_*K*_ is a positive semi-definite matrix factorised as **Σ**_*K*_ = **DCD**, where **D** = diag(*h*_1_, …, *h*_*K*_) and **C** *∝* |**C**|^*η*−1^ is a correlation matrix following the LKJ prior[32] with concentration parameter *η*. A value of *η* = 1 implies a uniform prior over correlation matrices, while *η* = 2 (the value we adopt here) concentrates more probability mass around the identity matrix. This structure for **Σ**_*K*_ implies both cluster-specific amplitudes 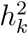, as well as correlations between clusters. Alternatively, we can assume that 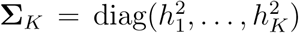, which implies that different clusters have different values of the amplitude parameters 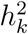, but are otherwise uncorrelated.

Finally, we examine the case where **Λ**_*MK×MK*_ is a block-diagonal matrix, with each of the *K* matrices along its main diagonal induced by the kernel 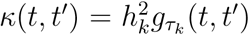, where both amplitude 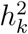 and time scale *τ*_*k*_ parameters are cluster-specific, but the clusters are otherwise uncorrelated.

### Relation between VAF and CCF

In this section, we give more details about the form of the function 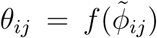, which encodes the relationship between VAF and CCF of mutation *i* in sample *j*. Each sample is viewed as a mixture of three cell populations[9]: a) a normal population of 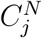 non-cancer cells, b) a reference population of 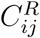 cancer cells, which do not harbour mutation *i* and c) a variant population of 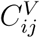 cancer cells, which harbour mutation *i*. The total number of cancer cells in the sample is 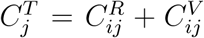. The reference and variant populations may each be further subdivided into sub-populations, where a different number of chromosomes covers locus *i* in each sub-population. The total number of chromosomes in the normal, reference and variant populations overlapping locus *i* in sample *j* are, respectively, equal to 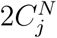 (assuming diploid normal cells), 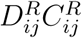 and 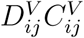, where 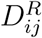 and 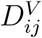 are the average numbers of chromosomes per cell covering locus *i* in sample *j* in each of the two cancer cell populations. Similarly, the total number of chromosomes harbouring mutation *i* in sample *j* is equal to 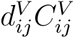, where 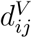 is the average number of chromosomes per cell harbouring mutation *i* in sample *j* in the variant cancer cell population (also known as *multiplicity*). We write:

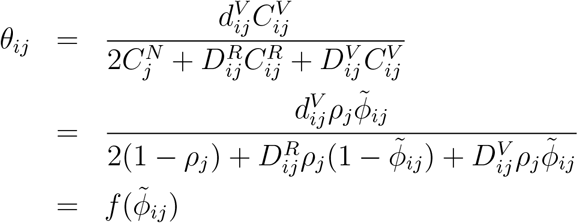

where 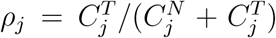 is the purity of the tumour and 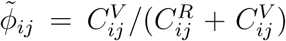. At this stage, two simplifying assumptions are often made: a) there are no subclonal copy number events, which implies that 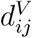, 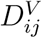 and 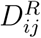 are integers, and b) the reference and variant cancer cell populations have the same copy number profile at locus *i* in sample *j*, i.e. 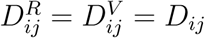. Under these assumptions, the above expression simplifies to:

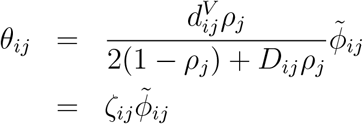

where *ζ*_*ij*_ is the value of *θ*_*ij*_ if mutation *i* in sample *j* is clonal (i.e. 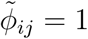). The quantities *ρ*_*j*_ and *D*_*ij*_ can be independently estimated (e.g. using software such as `ASCAT`[33], `ABSOLUTE`[34], `TITAN`[35] and others) and they are considered fixed. One way to approximate the multiplicity 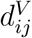 is as follows: first, we calculate 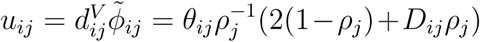. Then, we estimate 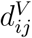 using the following rule:

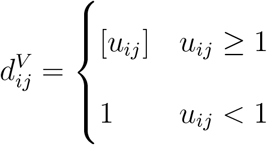

where [*u*_*ij*_] is the closest integer to *u*_*ij*_. For a justification of this estimation procedure, see [4].

### Observation models

We complete the above models by introducing expressions for the distribution of the read counts *r*_*ij*_ harbouring mutation *i* in sample *j*. Since high-throughput sequencing data often exhibit over-dispersion, we consider a beta-binomial model:

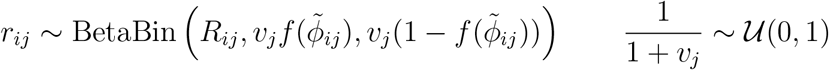

where *R*_*ij*_ is the sum of reads harbouring the alternative and reference alleles at locus *i* in sample *j* and 1/*v*_*j*_ is a sample-specific dispersion parameter. In the absence of over-dispersion (i.e. when *v*_*j*_ → ∞), the above reduces to the binomial model, 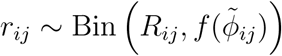.

### Inference

The above models were implemented using the Python-based probabilistic programming language `PyMC3 v3.8`[36] and inference was conducted using Automatic Differentiation Variational Inference (ADVI; [37]), instead of developing bespoke estimation algorithms, which is a rather laborious process particularly when multiple candidate models are considered [38–40]. Variational inference (VI; [41, 42]) is a computationally efficient approach for Bayesian inference, which aims to approximate the posterior density p(**z***|***y**) of latent variables **z** given data **y** using a surrogate probability density q_***θ***_(**z**) parametrised by a vector of variational parameters ***θ***. In our case, the data **y** are the locus- and sample-specific read counts *r*_*ij*_ and *r*_*ij*_, the local copy numbers *D*_*ij*_, the sample-specific purities *ρ*_*j*_ and the sample collection times *t*_*j*_, while the latent variables **z** are the cancer cell fractions *ϕ*_*jk*_, the cluster weights *w*_*k*_, the amplitudes 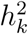, the time-scales *τ*_*k*_ and the sample-specific dispersions *v*_*j*_. VI approximates p(**z***|***y**) by maximising the lower bound of the marginal likelihood (or *evidence*) p(**y**), which is known as the *evidence lower bound* (ELBO), with respect to the variational parameters ***θ***:

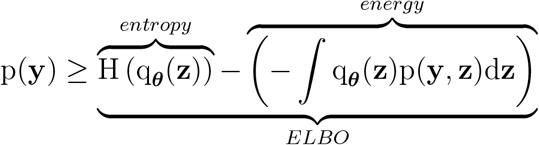

Maximising the ELBO is equivalent to jointly maximising the entropy term (which leads to a more spread out variational distribution q and prevents overfitting) and minimising the average energy term (i.e. the discrepancy between q and p). Furthermore, the maximised ELBO, being a lower bound of the evidence p(**y**), can be used for model comparison (see below).

### Performance metrics

We fit the above models against actual or simulated tumour samples (see Results). In the case of actual data, the ground truth (i.e. the actual clonal structure of each tumour sample) is unknown. In this case, we compare the performance of different models using the maximised ELBO (with a higher value indicating a better model). In the case of simulated data, the ground truth is known *a priori* and different models are compared using the *Adjusted Rand Index* (ARI), as implemented in the Python package `scikit-learn v0.22`[43]. ARI takes values between −1 and 1, with a value close to 1 or −1 indicating close agreement or disagreement to the ground truth, respectively, while a value close to 0 indicates random assignment of mutations to clusters. ARI is symmetric and for this reason we also use it for estimating the concordance between any two clustering models.

### Model nomenclature

In the Results section, the various models described above are referred to as follows. The model that assumes a uniform (i.e flat) prior over the CCF variables *ϕ*_*jk*_ is the **Flat** model. The model that assumes a single-output Gaussian Process prior over the transformed CCF variables *ψ*_*jk*_ is the **GP0** model. The models assuming a multi-output Gaussian Process prior on *ψ*_*jk*_ are labelled **GP1** (when **Σ**_*K*_ is diagonal), **GP2** (when **Σ**_*K*_ is full rank) and **GP3** (when **Λ**_*MK×MK*_ is block-diagonal with cluster-specific 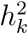 and *τ*_*k*_ parameters), respectively. Each of the models **GP0** to **GP3** admits exponential (**Exp**), **Mat32**, **Mat52** or exponentiated quadratic (**ExpQ**) kernels and are labelled accordingly, e.g. **GP0-Exp**, **GP0-ExpQ**, etc. In total, we examined 17 models. If the number of parameters in the **Flat** model is *n*_*p*_, the number of parameters in the **GP0**, **GP1**, **GP2** and **GP3** models is *n*_*p*_ + 2, *n*_*p*_ + *K* + 1, *n*_*p*_ + *K* + 1 + *K*(*K −* 1)*/*2 and *n*_*p*_ + 2*K*, respectively.

## RESULTS

We conducted a series of computational experiments on WES and WGS data from patients with CLL[23, 24], on TGS data from the liquid biopsy of a patient with melanoma[25] and on simulated data. The aim of these experiments was to demonstrate the application of the above models on longitudinal data and to assess their relative performance.

### The case of patient CLL003

First, we demonstrate the application of model **GP0-Mat32** on WGS data from patient CLL003 reported in [23] (Fig.1; the other models on the same dataset are shown in Fig.2). Details on sequencing and bioinformatics analysis for obtaining this data are given in the original paper. Briefly, peripheral blood was collected at five specific time points during disease progression, treatment and relapse together with a matched buccal swab (for germinal DNA). All samples underwent whole genome sequencing (WGS) followed by bioinformatics analysis, which identified 28 somatic mutations. Fitting the model to this data was performed by maximising the ELBO (see Methods), which can be used for assessing convergence of the estimation algorithm (typically achieved in less than 3K iterations; Fig.1A). Following a non-parametric approach for clustering mutations using a Dirichlet Process prior on the cancer cell fractions (see Methods) means that the number of clusters is not selected *a priori*, but rather estimated along with other model parameters (Fig.1B). We identified three major mutation clusters: one with median weight *≈*35% (i.e. any mutation has approximately 35% probability of belonging to this cluster) and two slightly smaller clusters with median *≈*30%. In Fig.1C, we illustrate the evolution of each cluster in time. Sample (a) was collected before commencing treatment with chlorambucil; sample (b) before treatment with fludarabine, cyclophosphamide and rituximab (FCR); sample (c) immediately after 6 cycles of FCR; sample (d) before treatment with ofatumumab; and sample (e) after treatment with ofatumumab, spanning in total a period of 35 months. Initial treatment with chlorambucil did not alter significantly the prevalence of the three mutation clusters, with median CCF>75% for clusters 1 and 3 and median CCF<10% for cluster 2. The second treatment regime (FCR) induced a dramatic reduction in the prevalence of cluster 3, but only a minor reduction of cluster 1. Concomitantly, the prevalence of cluster 2 increased substantially. By the end of the 35-months period, cluster 1 had recovered and, along with cluster 2, it reached CCF values of *≈*100%, while cluster 3 collapsed. Our algorithm soft-clusters mutations, i.e. for each mutation, it calculates the probability of membership to each cluster. From these, a hard clustering can be obtained by assigning each mutation to the cluster with the highest median membership probability. Fig.1D illustrates the hard cluster assignment for each mutation in the CLL003 dataset. It is interesting to observe that, by considering multiple time-separated samples, our method manages to deconvolve mutation clusters with similar VAF values, which would otherwise be hard to distinguish (e.g. observe the mixing of clusters 1 and 3 at timepoints (a) and (b) or clusters 1 and 2 at timepoints (d) and (e)). Finally, we can visually confirm the goodness of fit of the model to the data by overlaying the posterior predictive distribution (red lines in Fig.1E) on the histograms of observed VAF values for each sample.

**Figure 1:**
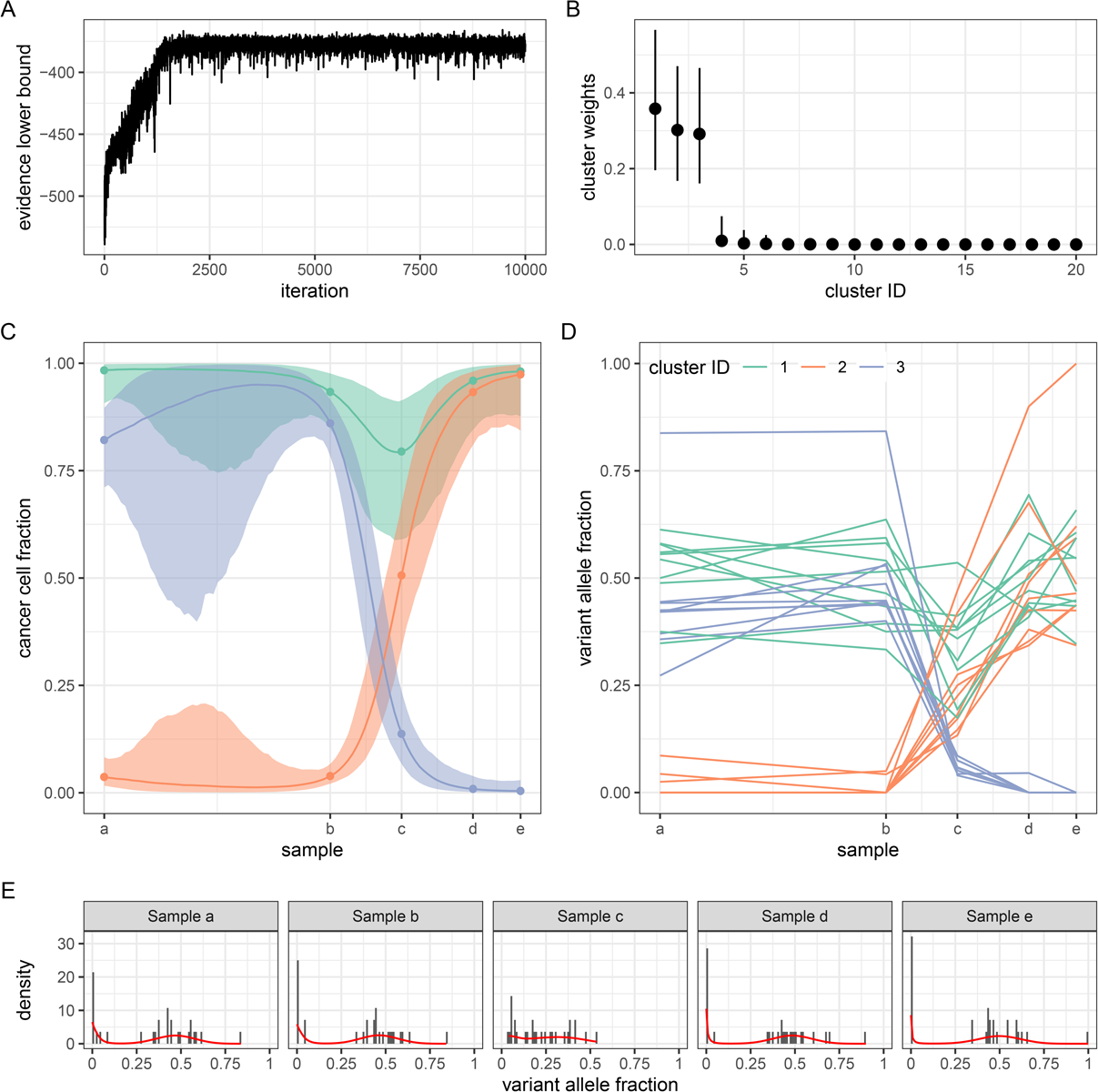
Application of model **GP0-Mat32** on data from patient CLL003[23]. A) Parameter estimation was achieved via maximisation of the evidence lower bound. Convergence was attained in less than 3K iterations. B) The number of clusters in the data was automatically estimated through the use of a Dirichlet Process prior. In this example, three major clusters were identified. C) The time profile of the three major clusters at each time point during disease treatment and progression. The median and 95% credible intervals are shown. Sample collection took place over the course of 35 motnhs. D) Observed VAF values for each somatic mutation and their cluster assignment. E) The fitted model (red lines) against the data in each sample.

**Figure 2:**
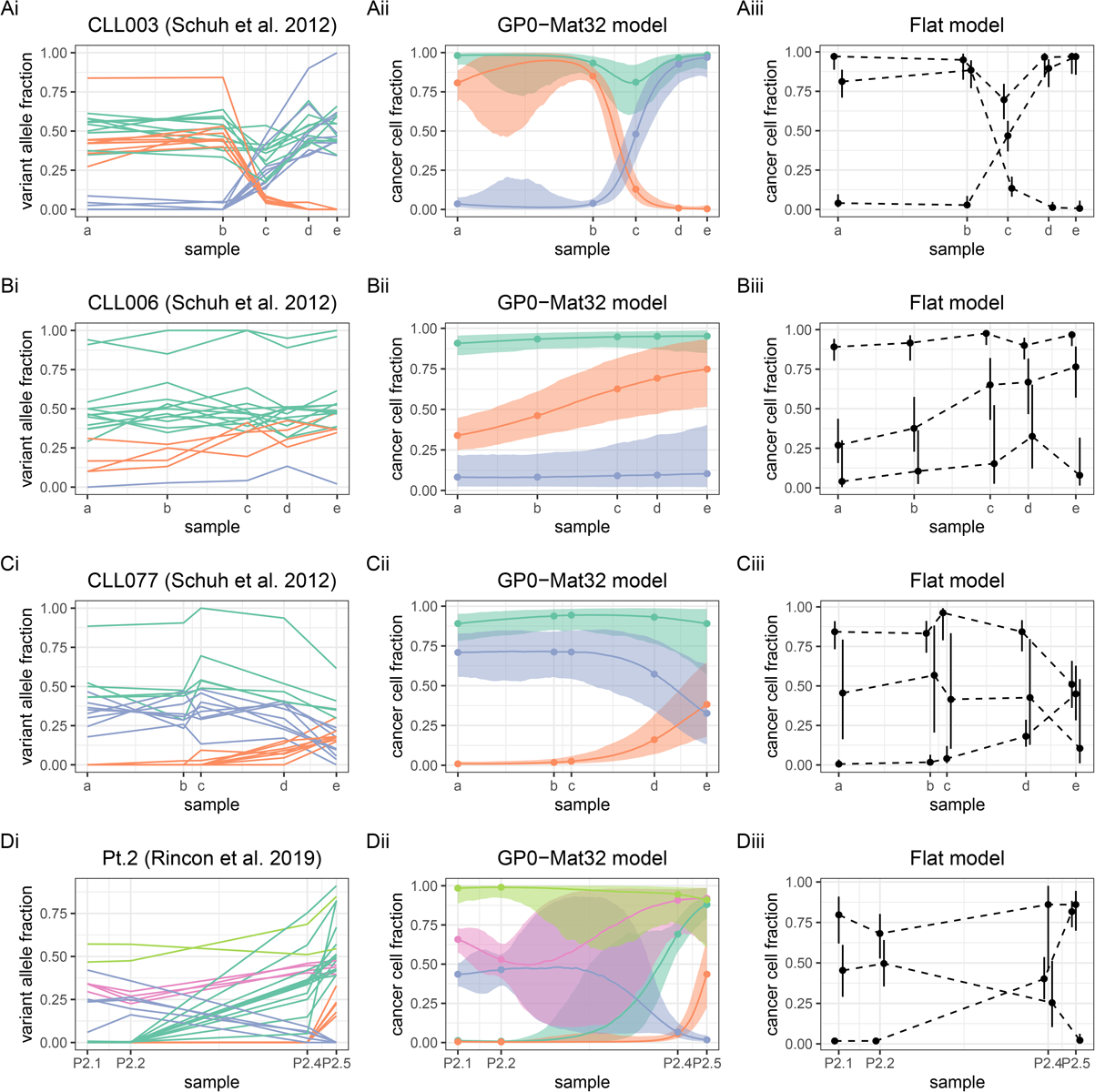
Overview of CLL data and fitted models **Flat** and **GP0-Mat32**. Unlike **GP0Mat32**, the **Flat** model estimates the cancer cell fraction of each cluster only at (but not between) the points of sample collection (dashed lines). Although both models identified the same number of cluster in datasets CLL003 to CLL077, the clusterings were not completely concordant (see main text for details).

### Benchmarks on CLL data with 4 or 5 samples

Next, we applied the remaining models on the data from patient CLL003, as well as all models on data from patients CL006 and CLL077 reported in [23] (Figs.2 and 3). WGS and bioinformatics analysis was conducted as for patient CLL003 (see original paper for details). For patient CLL006, sample (a) was collected before treatment with fludarabine and cyclophosphamide; sample (b) before treatment with rituximab; sample (c) before treatment with ofatumumab; sample (d) immediately after treatment with ofatumumab; and sample (e) at 12 months after treatment with ofatumumab spanning a period of 50 months. Similarly, for patient CLL077, sample (a) was collected before treatment with chlorambucil; sample (b) before treatment with fludarabine and cyclophosphamide; sample (c) immediately after 4 cycles of such treatment; sample (d) before treatment with ofatumumab; and sample (e) 9 months after ofatumumab spanning 57 months in total. In addition, we examined WES data from Patient 2 reported in [24]. Three peripheral blood mononulcear cell samples (P2.1, P2.2, P2.4) and one lymph node sample (P2.5) were collected over a period of 79 months, as well as a buccal mucosa sample for germinal DNA. Sample P2.1 was collected at diagnosis, sample P2.2 before treatment with FCR, sample P2.4 before treatment with TRU-016 and bendamustine and sample P2.4 before salvage chemotherapy and treatment with bortezomib (for details of sequencing and bioinformatics analysis, see original paper).

**Figure 3:**
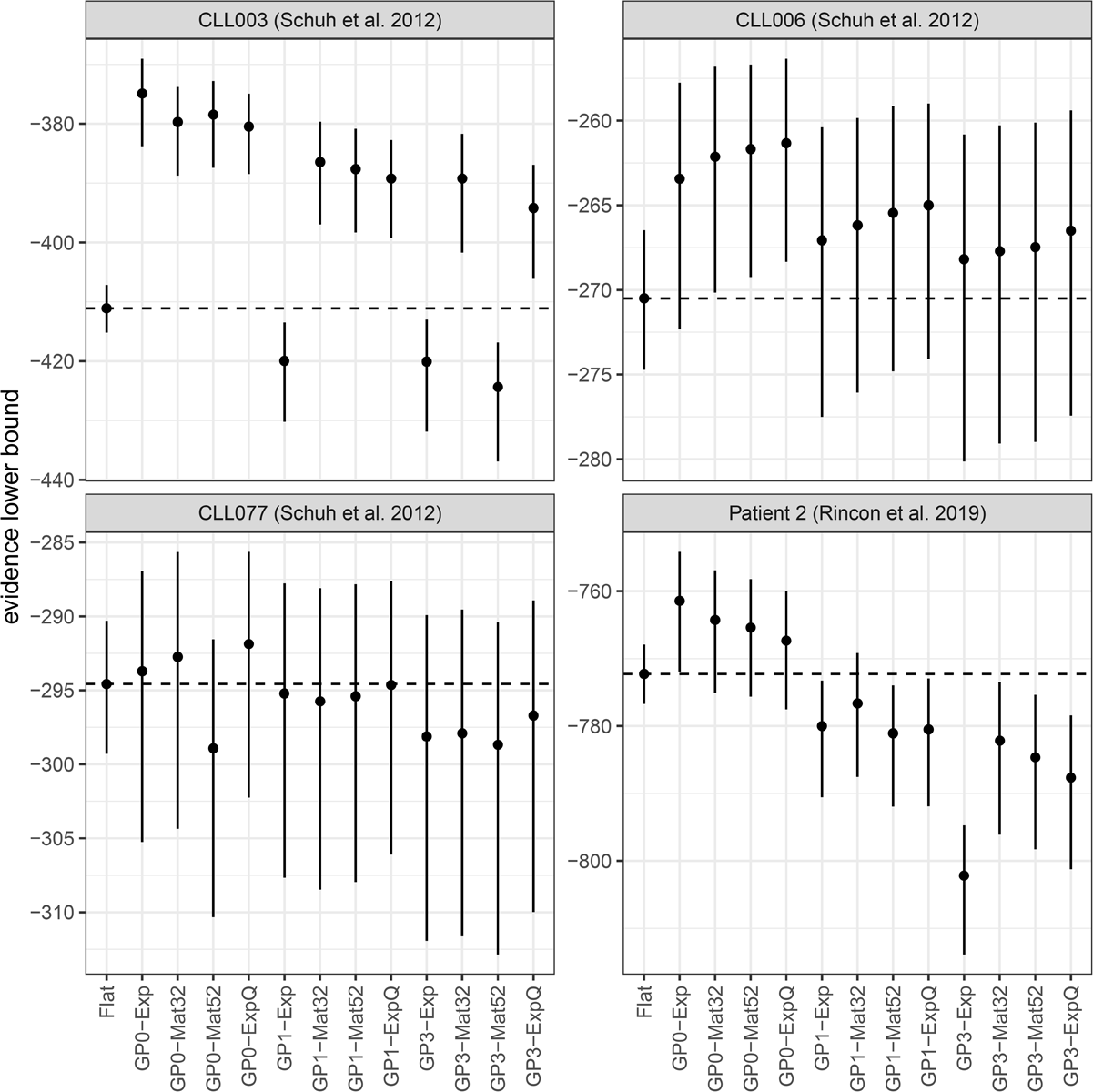
Comparison of all models using the data in the previous figure and the evidence lower bound for assessing performance. Members of the **GP0** group of models perform better than or comparably to the **Flat** model in all cases. Models **GP2** had the worst performance of all (as well as the largest number of parameters) and they were omitted from the figure.

There were 18, 21 and 32 somatic mutations in patients CLL006, CLL077 and Patient 2, respectively (as well as 28 somatic mutations in patient CLL003, as previously mentioned; Fig.2Ai-Di). A preliminary comparison indicates that, for patients CLL003 to CLL077, model **GP0-Mat32** (Figs. 3Aii-Cii) identified the same number of mutation clusters as the simpler **Flat** model (Fig.2Aiii-Ciii), i.e. three clusters with similar temporal dynamics. In order to assess the clustering concordance between the two models (i.e. whether they assign the same mutations to the same clusters), we calculated the values of ARI, which were equal to 0.54, 0.79 and 0.58, respectively. This indicates that the two models are not perfectly concordant in any of these three datasets (despite both identifying the same number of clusters) presumably due to the partial overlap between different mutation groups, as illustrated in Fig.2Ai-Ci. One striking difference between the **Flat** and **GP**-based models is that while the former estimates the latent state of the tumour only at the timepoints of sample collection (this is indicated by the dashed connecting lines in Fig.2Aiii-Ciii), the latter provides an estimate of the complete history of this latent state, i.e. both at and between these fixed timepoints. This is a major difference in favour of the use of **GP**-based models. In the case of Patient 2, the **Flat** and **GP0-Mat32** models identify three and five clusters, respectively (ARI=0.63; Fig.2Di-iii). For comparison, in the original paper, the authors identified seven clusters using `PyClone`[9].

In order to further assess the relative performance of different models (and without knowledge of the true clonal state of each tumour), we used the ELBO as performance metric (see Methods). The ELBO provides a lower bound on the marginal likelihood of the data (i.e. the evidence) and, at the same time, it includes an internal mechanism that prevents overfitting. Thus, it is often used in practise for model comparison and selection, with higher ELBO values indicating a better model. As illustrated in Fig.3A, all **GP0** models, all but one **GP1** models and all but two **GP3** models outperform the **Flat** model on the CLL003 data. The **GP2** model, which has by far the largest number of parameters, was the worst performer and it is omitted from the figure. There is a clear trend of decreasing performance with increasing number of parameters among the **GP**-based models, which is not surprising given that the lower the number of timepoints, the lower the capacity of the data to support overly complex models (as, for example, in the case of **GP2** models). In the case of CLL006 (Fig.3B), the same trend is observed, although the difference of the **GP**-based models from the **Flat** model is less pronounced. In the case of CLL077 (Fig.3C), models **GP0-Mat32** and **GP0-ExpQ** perform better than the **Flat** model (although this difference is not particularly pronounced because of the high variance of the ELBO), but the remaining **GP**-based models perform either clearly worse or comparably to the **Flat** model. In the case of Patient 2 (Fig.3D), the **GP0** models are again the best performers, unlike **GP1** and **GP3** models, which are clearly worse than the **Flat** model. In summary, there is always a member of the relatively parsimonious (in terms of the number of model parameters**) GP0** family of models that performs better than the **Flat** model in the above benchmarks.

### Benchmarks on CLL and melanoma data with 10 or 13 samples

Subsequently, we tested our models on longitudinal genomic data involving a higher number of timepoints. The first dataset comes from Patient 1 in [24]. A total of 13 peripheral blood mononuclear cell samples (P1.1 to P1.13) were collected over the course of 6.5 years and underwent targeted sequencing (TGS). Samples were collected before or after treatment commenced. In particular, sample P1.1 was collected before the patient received a stem cell transplant and the same holds for sample P1.8. Germinal DNA was obtained from a buccal mucosa sample and bioinformatics analysis identified 46 somatic mutations over all 13 samples (Fig.4A; see original paper for details). Model **GP0-Mat32** identified nine mutation clusters (Fig.4B), while the **Flat** model identified five (Fig.4C). In comparison, in the original paper, the authors estimated four clusters using PyClone[9]. Overall, models **GP0**, **GP1** and **GP2** perform better than the **Flat** model, unless an exponentiated quadratic kernel (**ExpQ**) is used (Fig.4D). We speculate that this is because **ExpQ** encodes perfectly smooth dynamics, which presumably cannot model sufficiently well the non-smooth bottleneck points P1.2 and P1.8 which precede stem cell transplantation. Model **GP3-Exp** is also performing better than the **Flat** model.

**Figure 4:**
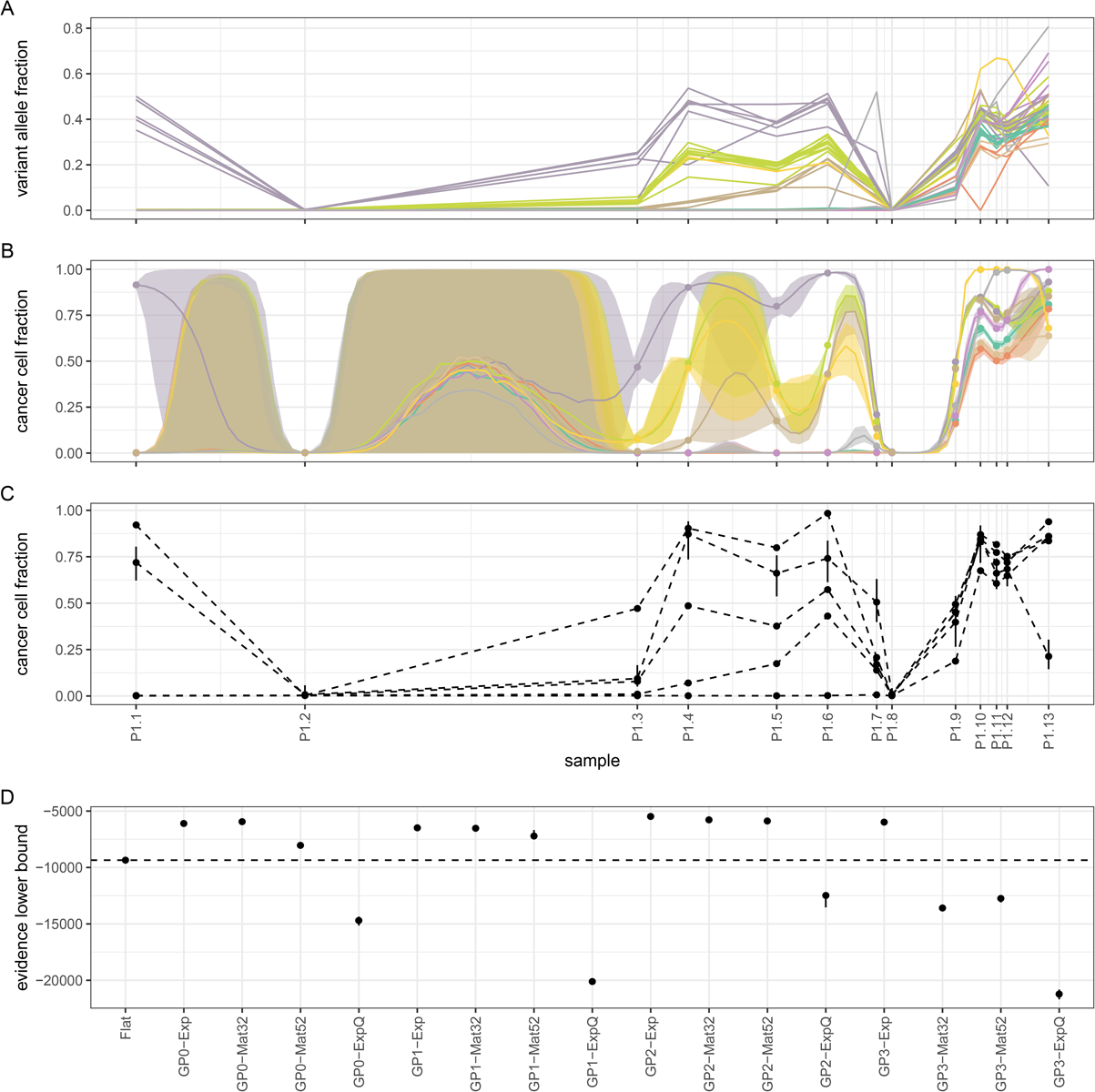
Assessing model performance on CLL data from Patient 1[24]. A) Observed VAF values for each somatic mutation over 6.5 years and their cluster assignments (colors are the same as in B). B) Mutation clusters identified by model **GP0-Mat32**. C) Mutation clusters identified by the **Flat** model. D) Comparative performance of various models. Notice that simpler models (**GP0**) often perform equivalently to or better than more complex ones (**GP1**, **GP2**, **GP3**).

The second multi-sample dataset comes from the liquid biopsy of a patient with metastatic melanoma[25]. Peripheral blood samples were collected at 10 different time points during pre-treatment, post-treatment and relapse over the course of 13 months. Germinal DNA was obtained from normal peripheral blood leucocytes. Targeted sequencing was conducted on extracted cell-free DNA followed by bioinformatics analysis, which revealed 63 somatic mutations. Visual inspection of the data indicates the absence of a definitive cluster structure (Fig.5A) and, for this reason, this is an interesting dataset to use for model evaluation. Both the **Flat** and **GP0-Exp** models identified five mutation clusters with little concordance between them (ARI=0.27) due to the extended overlap between different mutations bundles (Fig.5B,C). The median performance of model **GP0-Exp** is nominally higher than the **Flat** model, although it is doubtful whether the difference is substantial due to the high variance of the ELBO (Fig.5D). The remaining **GP**-based models perform worse than either **Flat** or **GP0-Exp**.

**Figure 5:**
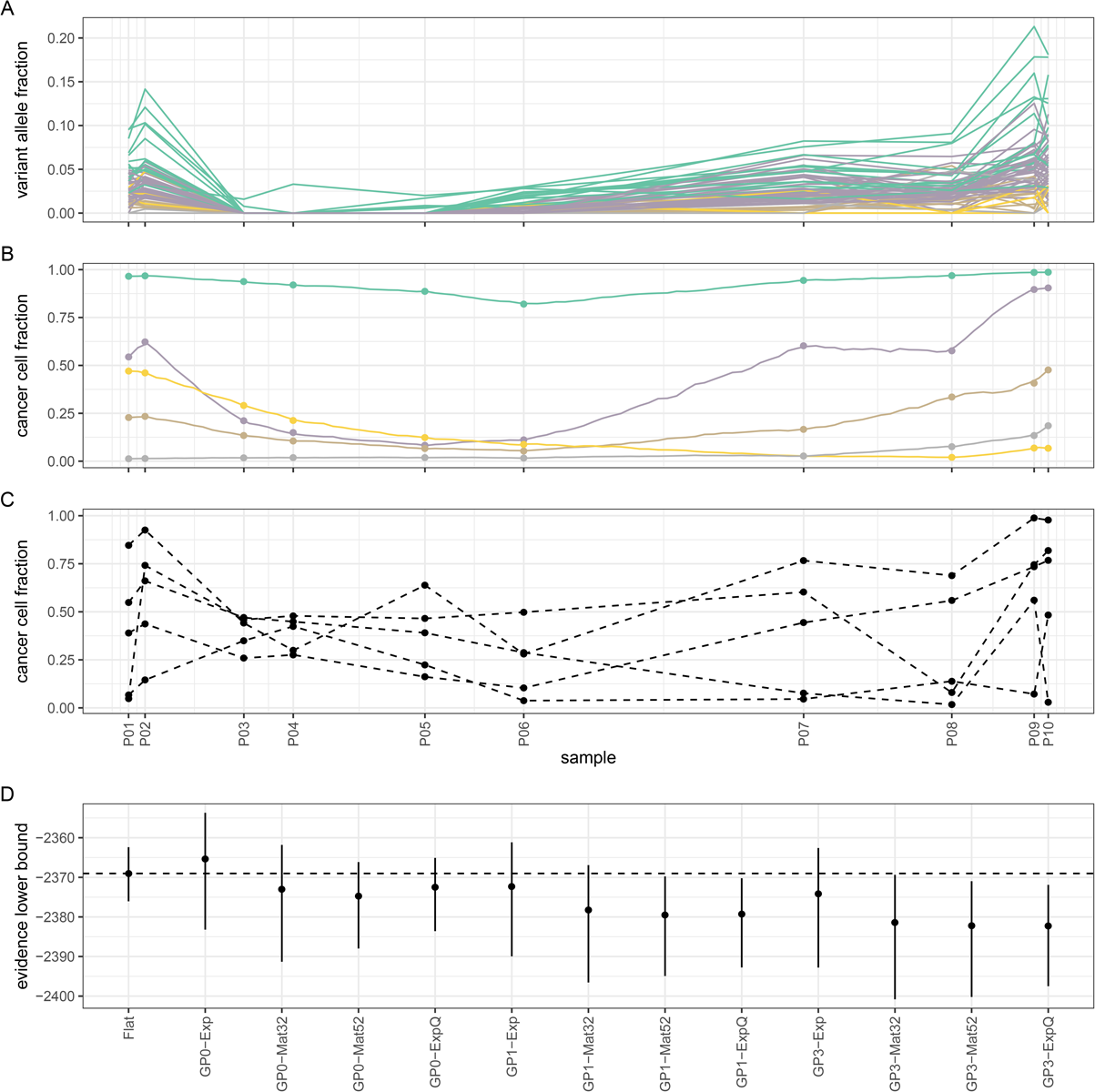
Assessing model performance using data from a liquid biopsy on a subject with melanoma[25]. A) Observed VAF values for each somatic mutation over 13 months of treatment and their cluster assignments (colors are the same as in B). B) Mutation clusters identified by model **GP0-Exp** (due to extensive overlap, credible intervals are omitted for clarity). C) Mutation clusters identified by the **Flat** model. D) Comparative performance of various models. Model **GP0-Exp** performs comparably to **Flat**.

### Computational experiments on simulated data

Overall, models **GP0** (particularly **GP0-Exp**) perform at least as well as the **Flat** model in all the above datasets. More complex models (i.e. models with a larger number of parameters), such as **GP1**, **GP2** and **GP3**, require a higher number of longitudinally collected samples for improved performance (Fig.4). However, this is not a sufficient condition, since data of low complexity (i.e. with trivial or non-obvious cluster structure and dynamics) can negatively affect the performance of the **GP**-based models (Fig.5).

We wanted to test whether these trends (i.e. the reduction in the performance of the **GP**-based models in relation to the **Flat** model as data size and complexity decreases) can be replicated using synthetic genomic data. For a given number of samples *M*, mutations *N* and mutation clusters *K*, data were simulated as follows (see source code on github for details): a) for each sample *j*, we randomly choose a purity value *ρ*_*j*_ between 80% and 90% and a random collection time *t*_*j*_ (with the first sample collected at time 0 and the last at time 1); b) for each cluster *k*, we sample a set of values *{ψ*_*jk*_}_*j,k*_ from a Gaussian process prior with squared amplitude *h*^2^ and inverse squared time scale *τ*; we calculate each *ϕ*_*jk*_ as a sigmoid function of *ψ*_*jk*_; c) for each mutation *i*, we randomly sample a cluster membership indicator *z*_*i*_ between 1 and *K*; d) finally, for each mutation *i* in each sample *j*, we sample the total number of reads *r*_*ij*_ from the empirical distribution of total reads in the data and then the number of mutated reads *r*_*ij*_ from a Binomial distribution: 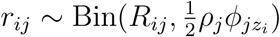. We generated data with *M* = {3, 6, 12}, *N* = {25, 50, 100} and *K* = {2, 4, 8}. For *h*^2^ and *τ*, we used the values {1, 10, 20} and {1, 10, 100}, respectively, which cover the range of values estimated from the actual data in the previous sections. For each of the 243 combinations of these parameters, we generated 3 replicates, which leads to a total of 729 datasets. Each such dataset was processed using the **Flat**, **GP0-Exp** and **GP0-Mat32** models (which were top performers on the actual data) and their performance was assessed against the true cluster structure of the dataset.

We may observe that when few samples are available (*M* = 3), the **Flat** model performs comparably to **GP0-Exp** and **GP0-Mat32** at all values of *N* and *K* (Fig.6). For medium (*M* = 6) and, particularly, large (*M* = 12) datasets, the **Flat** model starts falling behind the other two models, when the number of clusters in the data is relatively high (*K* = 4 or 8). These results indicate that in the presence of non-trivial cluster dynamics, the **Flat** model is comparable to **GP0-Exp** and **GP0-Mat32**, but only when the number of samples or data complexity (here, the number of clusters) is low.

**Figure 6:**
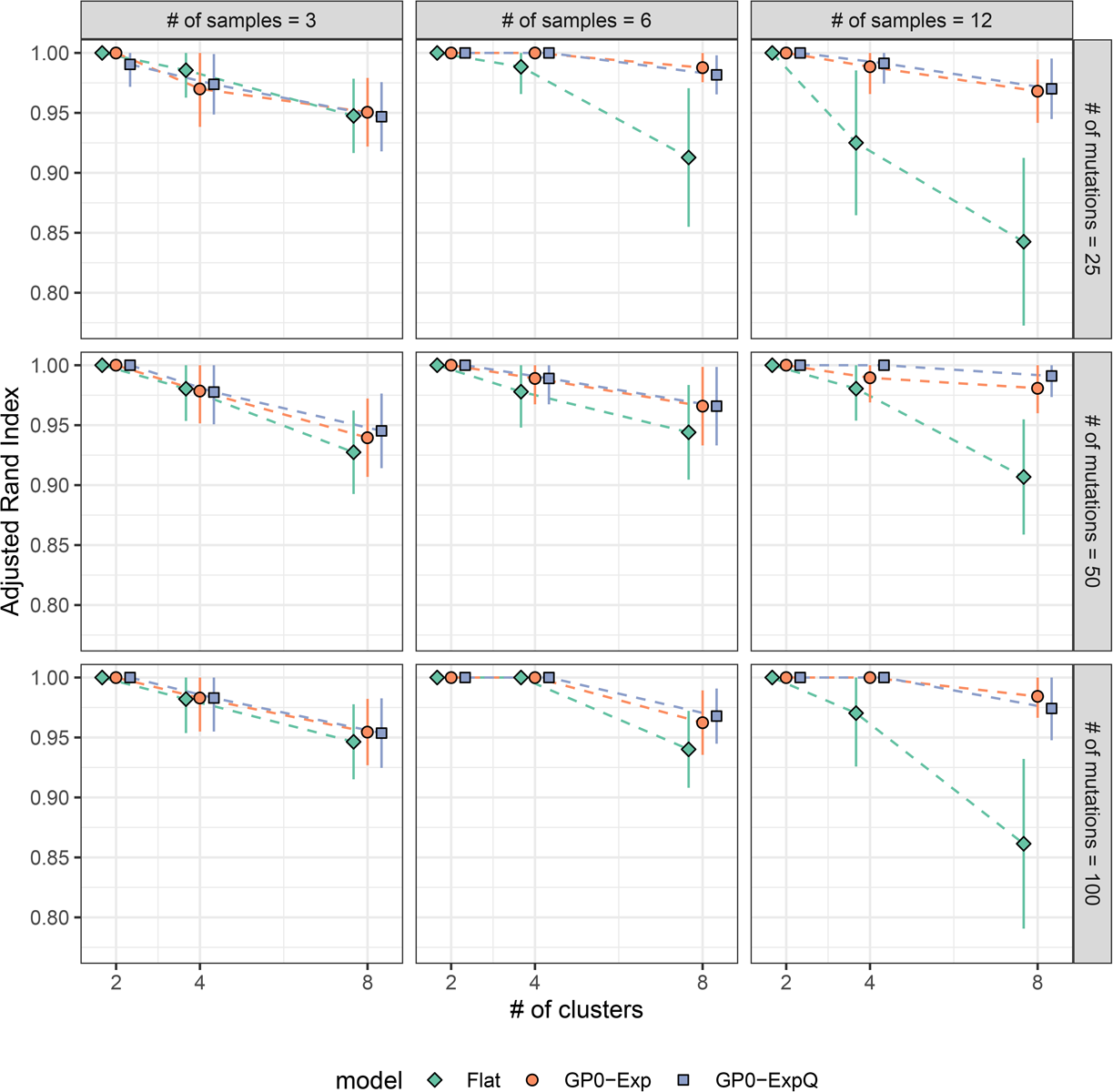
Benchmarks on synthetic data. When the number of samples is small (*M* = 3) or data complexity (i.e. the number of clusters) is low (*K* = 2), the **Flat** model performs comparably to the two **GP0** models. In all other cases, it is outperformed by them. Parameters used in data simulation were informed by the experimental data (see main text for details).

## DISCUSSION

Tumour heterogeneity in the form of distinct cancer cell populations or clones is the outcome of a process of continuous adaptation of the component cells to their local microenvironment. The outcome of any therapeutic intervention depends on this latent cellular diversity and, for this reason, statistical methodologies that help deconvolve the clonal structure of tumours are valuable tools at the disposal of clinicians and bioinformaticians.

In this paper, we proposed a statistical methodology for clonal deconvolution based on longitudinal data, which explicitly takes into account the temporal spacing of sample collection. Our approach combines two Bayesian non-parametric statistical frameworks, namely Dirichlet Process Mixture Models (for clustering in the absence of prior knowledge on the number of clusters supported by the data) and Gaussian Process Latent Variable Models (for modelling the time-dependence of clone prevalence without any explicit assumptions on the form of this dependence). Using a combination of experimental data from patients with CLL or melanoma, as well as synthetic data simulated using experimental data as template, we demonstrate that there are advantages in this approach, as long as data of sufficient volume and complexity are available. When this is not the case, our methodology still manages to reconstruct the time dependence of mutation clusters continuously in time (i.e. at and between sampling timepoints) from a small number of sequentially collected samples.

CLL is an ideal experimental model for the study of cancer evolution, because it develops over many years and because the collection of a long sequence of blood samples from the same patient for genomic analysis is easy, at least when compared to solid tumours. Thus, we expect that our methodology will find applications in the study of CLL and other liquid cancers. It can also be used as a general purpose clustering tool for identifiying populations of mutations based on sequencing of circulating tumour DNA obtained through a liquid biopsy.

As with other approaches for clustering mutations based on bulk sequencing data, a phylogeny is not derived directly, but it can be calculated retrospectively using the output of our method as input to bespoke software[44–46]. Furthemore, single-cell sequencing promises to alleviate the confounding of clones inherent in methods based on bulk sequencing by permitting direct observation of the genotypes of the cells that compose each clone. However, it is in turn plaqued by its own technical limitations, namely high levels of noise, error rates and missing values[47–56].

In conclusion, we propose that taking into account information on the temporal spacing of longitudinal tumour samples can impove clonal deconvolution and we show how this can be achieved in the context of non-parametric Bayesian statistics.

## REFERENCES

[1] P C Nowell. ‘The clonal evolution of tumor cell populations’. en. In: Science 194.4260 (Oct. 1976), pp. 23–28.

[2] Lauren M F Merlo et al. ‘Cancer as an evolutionary and ecological process’. en. In: Nat. Rev. Cancer 6.12 (Dec. 2006), pp. 924–935.

[3] Niko Beerenwinkel et al. ‘Cancer evolution: mathematical models and computational inference’. en. In: Syst. Biol. 64.1 (Jan. 2015), e1–25.

[4] Stefan C Dentro, David C Wedge and Peter Van Loo. ‘Principles of Reconstructing the Subclonal Architecture of Cancers’. en. In: Cold Spring Harb. Perspect. Med. 7.8 (Aug. 2017).

[5] Wazim Mohammed Ismail, Etienne Nzabarushimana and Haixu Tang. ‘Algorithmic approaches to clonal reconstruction in heterogeneous cell populations’. In: Quantitative Biology 7.4 (Dec. 2019), pp. 255–265.

[6] Adriana Salcedo et al. ‘A community effort to create standards for evaluating tumor subclonal reconstruction’. en. In: Nat. Biotechnol. 38.1 (Jan. 2020), pp. 97–107.

[7] Andrej Fischer et al. ‘High-definition reconstruction of clonal composition in cancer’. en. In: Cell Rep. 7.5 (June 2014), pp. 1740–1752.

[8] Habil Zare et al. ‘Inferring clonal composition from multiple sections of a breast cancer’. en. In: PLoS Comput. Biol. 10.7 (July 2014), e1003703.

[9] Andrew Roth et al. ‘PyClone: statistical inference of clonal population structure in cancer’. en. In: Nat. Methods 11.4 (Apr. 2014), pp. 396–398.

[10] Subhajit Sengupta et al. ‘BayClone: Bayesian non-parametric inference of tumour sub-clones using NGS data’. In: Biocomputing 2015. WORLD SCIENTIFIC, Nov. 2014, pp. 467–478.

[11] Christopher A Miller et al. ‘SciClone: inferring clonal architecture and tracking the spatial and temporal patterns of tumor evolution’. en. In: PLoS Comput. Biol. 10.8 (Aug. 2014), e1003665.

[12] Amit G Deshwar et al. ‘PhyloWGS: reconstructing subclonal composition and evolution from whole-genome sequencing of tumors’. en. In: Genome Biol. 16 (Feb. 2015), p. 35.

[13] Ke Yuan et al. ‘BitPhylogeny: a probabilistic framework for reconstructing intra-tumor phylogenies’. en. In: Genome Biol. 16 (Feb. 2015), p. 36.

[14] Mohammed El-Kebir et al. ‘Reconstruction of clonal trees and tumor composition from multi-sample sequencing data’. en. In: Bioinformatics 31.12 (June 2015), pp. i62–70.

[15] Yuchao Jiang et al. ‘Assessing intratumor heterogeneity and tracking longitudinal and spatial clonal evolutionary history by next-generation sequencing’. en. In: Proc. Natl. Acad. Sci. U. S. A. 113.37 (Sept. 2016), E5528–37.

[16] Francesco Marass et al. ‘A phylogenetic latent feature model for clonal deconvolution’. en. In: Ann. Appl. Stat. 10.4 (Dec. 2016), pp. 2377–2404.

[17] Nilgun Donmez et al. ‘Clonality Inference from Single Tumor Samples Using Low-Coverage Sequence Data’. en. In: J. Comput. Biol. 24.6 (June 2017), pp. 515–523.

[18] Yulia Rubanova et al. ‘TrackSig: reconstructing evolutionary trajectories of mutations in cancer’. en. Nov. 2018.

[19] Ke Yuan et al. ‘Ccube: A fast and robust method for estimating cancer cell fractions’. en. Dec. 2018.

[20] Matthew A Myers, Gryte Satas and Benjamin J Raphael. ‘CALDER: Inferring Phylogenetic Trees from Longitudinal Tumor Samples’. en. In: Cell Syst 8.6 (June 2019), 514–522.e5.

[21] Judith Abécassis, Fabien Reyal and Jean-Philippe Vert. ‘CloneSig: Joint inference of intra-tumor heterogeneity and signature deconvolution in tumor bulk sequencing data’. en. Oct. 2019.

[22] Mark R Zucker et al. ‘Inferring Clonal Heterogeneity in Cancer using SNP Arrays and Whole Genome Sequencing’. en. In: Bioinformatics (Jan. 2019).

[23] Anna Schuh et al. ‘Monitoring chronic lymphocytic leukemia progression by whole genome sequencing reveals heterogeneous clonal evolution patterns’. en. In: Blood 120.20 (Nov. 2012), pp. 4191–4196.

[24] Julia González-Rincón et al. ‘Clonal dynamics monitoring during clinical evolution in chronic lymphocytic leukaemia’. en. In: Sci. Rep. 9.1 (Jan. 2019), p. 975.

[25] Anthony Cutts et al. ‘Characterisation of the changing genomic landscape of metastatic melanoma using cell free DNA’. en. In: NPJ Genom Med 2 (Sept. 2017), p. 25.

[26] Andrew Gelman et al. Bayesian Data Analysis (Chapman & Hall/CRC Texts in Statistical Science). en. 3 edition. Chapman and Hall/CRC, Nov. 2013.

[27] Serena Nik-Zainal et al. ‘The life history of 21 breast cancers’. en. In: Cell 149.5 (May 2012), pp. 994–1007.

[28] Niccolo Bolli et al. ‘Heterogeneity of genomic evolution and mutational profiles in multiple myeloma’. en. In: Nat. Commun. 5 (2014), p. 2997.

[29] Carl Edward Rasmussen and Christopher K I Williams. Gaussian Processes for Machine Learning. en. MIT Press, Jan. 2006.

[30] Roberts S. et al. ‘Gaussian processes for time-series modelling’. In: Philosophical Transactions of the Royal Society A: Mathematical, Physical and Engineering Sciences 371.1984 (Feb. 2013), p. 20110550.

[31] Mauricio A Álvarez, Lorenzo Rosasco and Neil D Lawrence. ‘Kernels for Vector-Valued Functions: A Review’. In: Foundations and Trends® in Machine Learning 4.3 (2012), pp. 195–266.

[32] Stan Development Team. 1.13 Multivariate Priors for Hierarchical Models | Stan User’s Guide. https://mc-stan.org/docs/2_21/stan-users-guide/multivariate-hierarchical-priors-section.html. Accessed: 2020-1-17.

[33] Peter Van Loo et al. ‘Allele-specific copy number analysis of tumors’. en. In: Proc. Natl. Acad. Sci. U. S. A. 107.39 (Sept. 2010), pp. 16910–16915.

[34] Scott L Carter et al. ‘Absolute quantification of somatic DNA alterations in human cancer’. en. In: Nat. Biotechnol. 30.5 (May 2012), pp. 413–421.

[35] Gavin Ha et al. ‘TITAN: inference of copy number architectures in clonal cell populations from tumor whole-genome sequence data’. en. In: Genome Res. 24.11 (Nov. 2014), pp. 1881–1893.

[36] John Salvatier, Thomas V Wiecki and Christopher Fonnesbeck. ‘Probabilistic programming in Python using PyMC3’. en. In: PeerJ Comput. Sci. 2 (Apr. 2016), e55.

[37] Alp Kucukelbir et al. ‘Automatic Differentiation Variational Inference’. In: J. Mach. Learn. Res. 18.14 (2017), pp. 1–45.

[38] Dimitrios V Vavoulis et al. ‘DGEclust: differential expression analysis of clustered count data’. en. In: Genome Biol. 16 (Feb. 2015), p. 39.

[39] Dimitrios V Vavoulis, Jenny C Taylor and Anna Schuh. ‘Hierarchical probabilistic models for multiple gene/variant associations based on next-generation sequencing data’. en. In: Bioinformatics 33.19 (Oct. 2017), pp. 3058–3064.

[40] Dimitrios V Vavoulis. ‘Exploring Bayesian Approaches to eQTL Mapping Through Probabilistic Programming’. en. In: Methods Mol. Biol. 2082 (2020), pp. 123–146.

[41] David M Blei, Alp Kucukelbir and Jon D McAuliffe. ‘Variational Inference: A Review for Statisticians’. In: J. Am. Stat. Assoc. 112.518 (Apr. 2017), pp. 859–877.

[42] Cheng Zhang et al. ‘Advances in Variational Inference’. en. In: IEEE Trans. Pattern Anal. Mach. Intell. (Dec. 2018).

[43] Fabian Pedregosa et al. ‘Scikit-learn: Machine Learning in Python’. In: J. Mach. Learn. Res. 12.Oct (2011), pp. 2825–2830.

[44] Yi Qiao et al. ‘SubcloneSeeker: a computational framework for reconstructing tumor clone structure for cancer variant interpretation and prioritization’. en. In: Genome Biol. 15.8 (Aug. 2014), p. 443.

[45] Noushin Niknafs et al. ‘SubClonal Hierarchy Inference from Somatic Mutations: Automatic Reconstruction of Cancer Evolutionary Trees from Multi-region Next Generation Sequencing’. en. In: PLoS Comput. Biol. 11.10 (Oct. 2015), e1004416.

[46] H X Dang et al. ‘ClonEvol: clonal ordering and visualization in cancer sequencing’. en. In: Ann. Oncol. 28.12 (Dec. 2017), pp. 3076–3082.

[47] Andrew Roth et al. ‘Clonal genotype and population structure inference from single-cell tumor sequencing’. en. In: Nat. Methods 13.7 (July 2016), pp. 573–576.

[48] Edith M Ross and Florian Markowetz. ‘OncoNEM: inferring tumor evolution from single-cell sequencing data’. en. In: Genome Biol. 17 (Apr. 2016), p. 69.

[49] Katharina Jahn, Jack Kuipers and Niko Beerenwinkel. ‘Tree inference for single-cell data’. en. In: Genome Biol. 17 (May 2016), p. 86.

[50] Hamim Zafar et al. ‘SiFit: inferring tumor trees from single-cell sequencing data under finite-sites models’. en. In: Genome Biol. 18.1 (Sept. 2017), p. 178.

[51] Mohammed El-Kebir. ‘SPhyR: tumor phylogeny estimation from single-cell sequencing data under loss and error’. en. In: Bioinformatics 34.17 (Sept. 2018), pp. i671–i679.

[52] Hamim Zafar et al. ‘SiCloneFit: Bayesian inference of population structure, genotype, and phylogeny of tumor clones from single-cell genome sequencing data’. en. In: Genome Res. 29.11 (Nov. 2019), pp. 1847–1859.

[53] Salem Malikic et al. ‘PhISCS: a combinatorial approach for subperfect tumor phylogeny reconstruction via integrative use of single-cell and bulk sequencing data’. en. In: Genome Res. 29.11 (Nov. 2019), pp. 1860–1877.

[54] Ziwei Chen et al. ‘RobustClone: A robust PCA method of tumor clone and evolution inference from single-cell sequencing data’. en. June 2019.

[55] Nico Borgsmueller et al. ‘Bayesian non-parametric clustering of single-cell mutation profiles’. en. Jan. 2020.

[56] Daniele Ramazzotti et al. ‘Longitudinal cancer evolution from single cells’. en. Jan. 2020.

